# Single-cell replication profiling reveals stochastic regulation of the mammalian replication-timing program

**DOI:** 10.1101/158352

**Authors:** Vishnu Dileep, David M. Gilbert

**Affiliations:** Department of Biological Science, 319 Stadium Drive, Florida State University, Tallahassee, FL 32306, USA.

## Abstract

In mammalian cells, distinct replication domains (RDs), corresponding to structural units of chromosomes called topologically-associating domains (TADs), replicate at different times during S-phase^1–4^. Further, early/late replication of RDs corresponds to active/inactive chromatin interaction compartments^5,6^. Although replication origins are selected stochastically, such that each cell is using a different cohort of origins to replicate their genomes^7–12^, replication-timing is regulated independently and upstream of origin selection^13^ and evidence suggests that replication timing is conserved in consecutive cell cycles^14^. Hence, quantifying the extent of cell-to-cell variation in replication timing is central to studies of chromosome structure and function. Here we devise a strategy to measure variation in single-cell replication timing using DNA copy number. We find that borders between replicated and un-replicated DNA are highly conserved between cells, demarcating active and inactive compartments of the nucleus. Nonetheless, measurable variation was evident. Surprisingly, we detected a similar degree of variation in replication timing from cell-to-cell, between homologues within cells, and between all domains genome-wide regardless of their replication timing. These results demonstrate that stochastic variation in replication timing is independent of elements that dictate timing or extrinsic environmental variation.

---

Single-cell DNA copy number can distinguish replicated DNA from un-replicated DNA^15,16^. Specifically, regions that have completed replication will have twice the copy number compared to regions that have not replicated. Hence, we reasoned that measurements of DNA copy number in cells isolated at different times during S-phase could reveal replication-timing programs in single-cells. Moreover, to separately evaluate the extent of extrinsic (cell-to-cell) vs. intrinsic (homologue-to-homologue) variability in replication timing, we examined both haploid H129-2 mouse embryonic stem cells (mESCs) and diploid hybrid *musculus* 129 × *Castaneus* mESCs that harbor a high single nucleotide polymorphism (SNP) density between homologues, permitting allele specific analysis. To generate single-cell copy number variation (CNV) profiles, we used flow cytometry of DNA-stained cells to sort single S-phase cells into 96 well plates followed by whole genome amplification (WGA). Amplified DNA from each cell was uniquely barcoded and sequenced (Fig. 1a)^17,18^. To control for amplification and mappability biases, we also sorted G1 and G2 cells, which contain a uniform DNA content. Regions of low mappability and over amplification were removed based on the G1 and G2 controls. Read counts were normalized by dividing the coverage data of each single-cell by the coverage of the G1 and G2 control cells. Next, a median filter was applied to smooth the data, producing CNV profiles in 50kb bins (Methods).

**Figure. 1:**
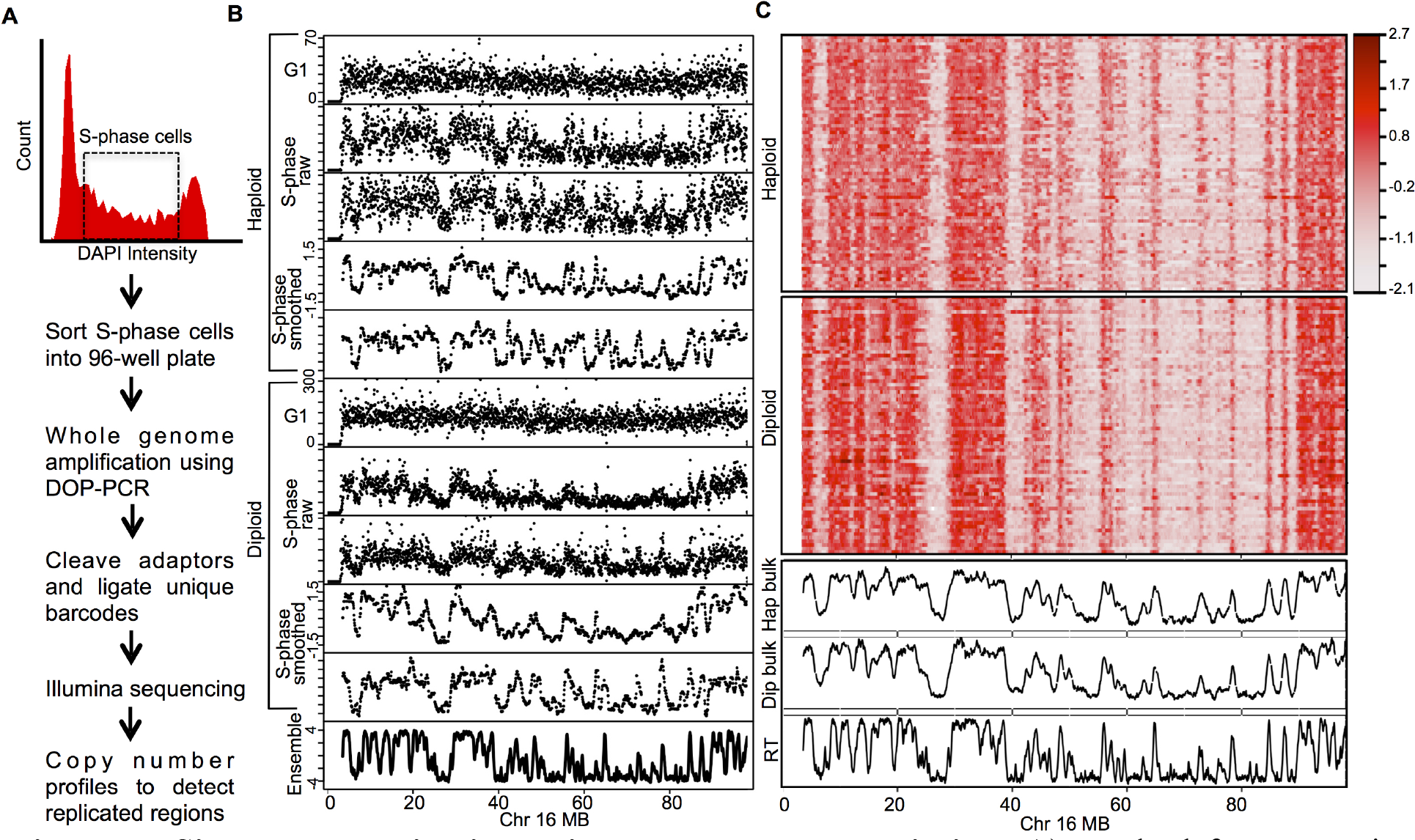
Single-cell replication using copy number variation. a) Method for generating single-cell CNV profiles b) Representative single-cell CNV profiles of G1 and S-phase cells in both haploid and diploid hybrid cells. CNV profiles are shown as raw read count in 50kb bins and after smoothing and corrections. c) Heatmap of all single-cell CNV profiles after smoothing and corrections. The bottom 3 panels show average of haploid single-cells, average of diploid single-cells and replication timing measured using population-based BrdU-IP in the diploid hybrid cells.

We generated single-cell sequencing data for 200 mESCs, composed of 93 haploid H129-2 mESCs and 107 129 × *Castaneus* diploid mESCs. Since we expected the CNV profile of mid-S-phase cells to distinguish the maximum number of early and late replicating domains, we sorted a majority of the cells from mid-S-phase. We extended the sorting gates for the haploid H129-2 cell line to early and late S-phase (Supplementary Fig. 1a). Cells with few reads (5.5% of cells) or cells with complex karyotype aberrations/complete loss of a chromosome (7% of cells) were discarded, while cells with aneuploidy were corrected by normalizing to the mean read density for those chromosomes in the control G1 and G2 cells (see Methods). Approximately, 27% of the diploid hybrid cells showed karyotype aberrations or aneuploidy, with the most frequent being X-chromosome loss, consistent with previous observations^19,20^. Only 6% of haploid cells showed karyotype aberrations or aneuploidy (Supplementary Fig. 2).

**Figure. 2:**
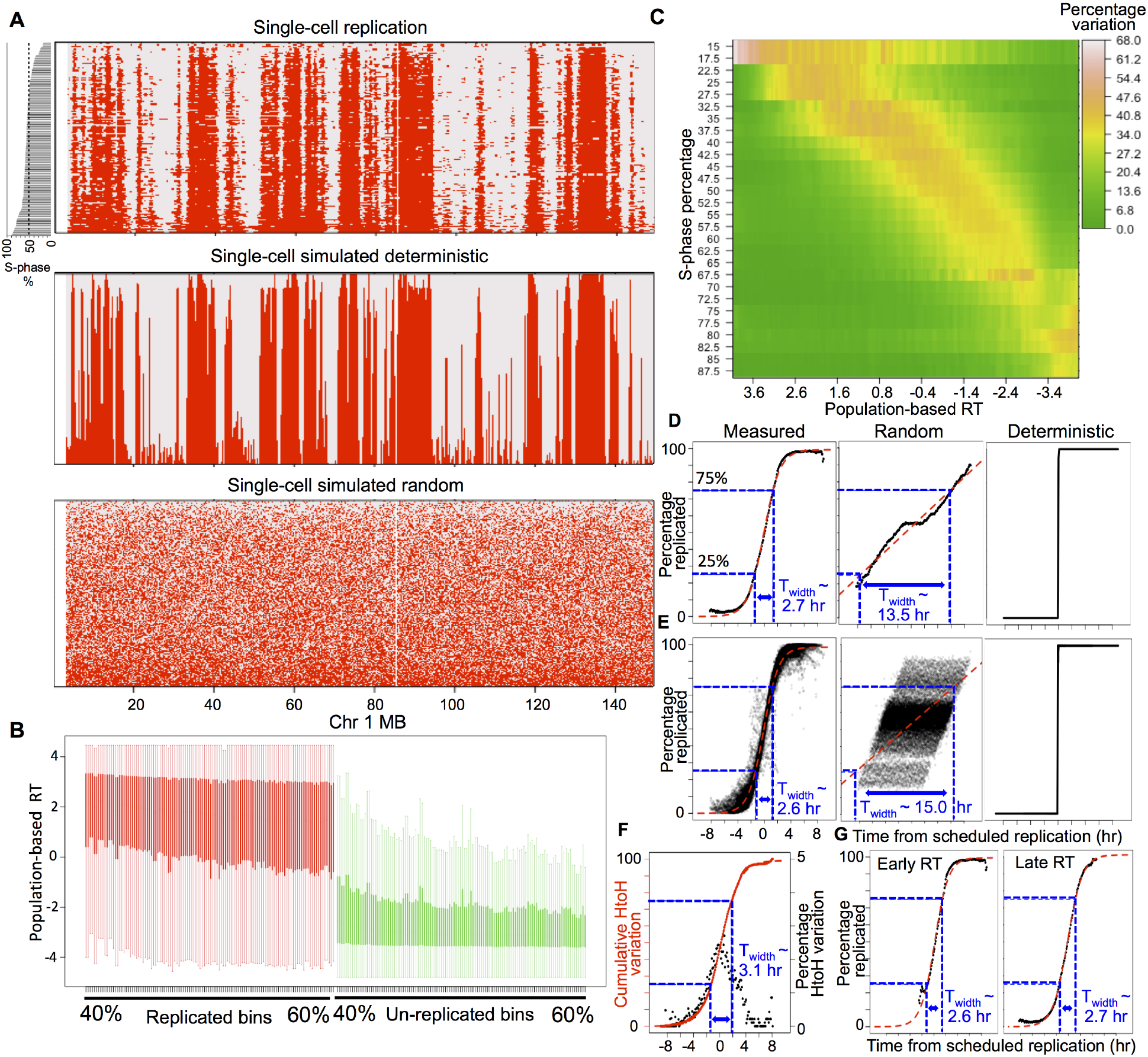
Measuring single-cell replication variability after binarization. a) Binarized replication status in all haploid single cells and homologue-parsed diploid cells. The cells are ranked by its progress in S-phase which is plotted as a bar plot on the left. The bottom panels show simulated deterministic and simulated random replication for identical S-phase distribution of single-cells (methods). b) Boxplot of population-based replication timing for replicated bins (red) and un-replicated bins (green) for each single cell ranked between 40 to 60 percentages of S-phase progression. c) Heatmap of variability between pairs of cells ranked within one percentile of each other. The Y-axis is average s-phase rank of pairs of cells and X-axis is the replication timing (RT) measured by population-based method. The Y-axis is measured in intervals of 5% of S-phase progression with a step size of 2.5 and the X-axis is measured is intervals of 0.1 population-based replication timing score with a step-size of 0.05. The pair-wise variability between cells is (measured using the binarized data) is the percentage of 50kb bins where there is a transition in the binary signal for the given S-phase progression interval and population-based RT interval. d) Cell-to-cell variability vs. “time from scheduled replication” hours. The mean across all 50kb bins is plotted for each 0.1hr interval on x-axis and the red line is the sigmoid fit. e) Within cell variability across the genome plotted for all single cells similar to panle D. f) Homologue-to-homologue variability vs “time from scheduled replication” (black). Solid red line is cumulative sum of the variability and the dotted red is the sigmoid fit. g) Cell-to-cell variability for early replicating bins (RT > 0) and late replicating bins (RT < 0).

Cells sorted from the middle of S-phase showed an oscillating CNV relative to the flat CNV profile for cells sorted from G1 or G2 phase (Fig. 1b). Regions with higher copy number aligned with early replicating domains detected by population-based replication timing data measured using Repli-seq^21^. Correcting the raw read counts for mappability and WGA biases, followed by smoothing and scaling the data, produced CNV profiles with reduced noise without any loss of information (Fig. 1b). Population-based replication timing profiles have revealed chromosomal segments that replicate relatively synchronously, appearing as a plateau in RT profiles. These segments are termed Constant Timing Regions (CTRs). Heat maps of single-cell replication-timing profiles from across all single-cells show clear signatures of domains that correspond to CTRs indicating conservation of replication timing at the single-cell level (Fig. 1c). Average replication-timing profiles generated from both the haploid and the diploid cells show a Pearson correlation of ∼0.89 with population-based Repli-seq profiles, demonstrating the robustness of our protocol and computational pipeline.

Haploid and diploid cells were processed differently. For haploid single-cells, replication data can assume 2 states; replicated or un-replicated, hence, we binarized the corrected data to reflect the replicated vs. un-replicated states as follows. First, we segmented the smoothed CNV profiles to identify segments with higher and lower copy number. Next, for each cell, we generated several binary signals using evenly spaced threshold values. Finally, we chose the threshold value at which the distance (Manhattan) between the binary signal and the un-binarized segmented data was minimum (Methods, **Supplementary Video 1**). Binary classification allowed us to rank individual cells within the spectrum of S-phase progression based on the number of bins identified as replicated. The ranking accurately reflected the early, middle or late FACS sorting gates used to collect the single-cells (Supplementary Fig. 1b). Outlier cells that did not correlate with any other cells were removed from further analysis (Supplementary Fig. 3, methods). At the end of these data processing steps, 75 haploid S-phase cells had passed all quality control measures and were used for further analysis.

**Figure. 3:**
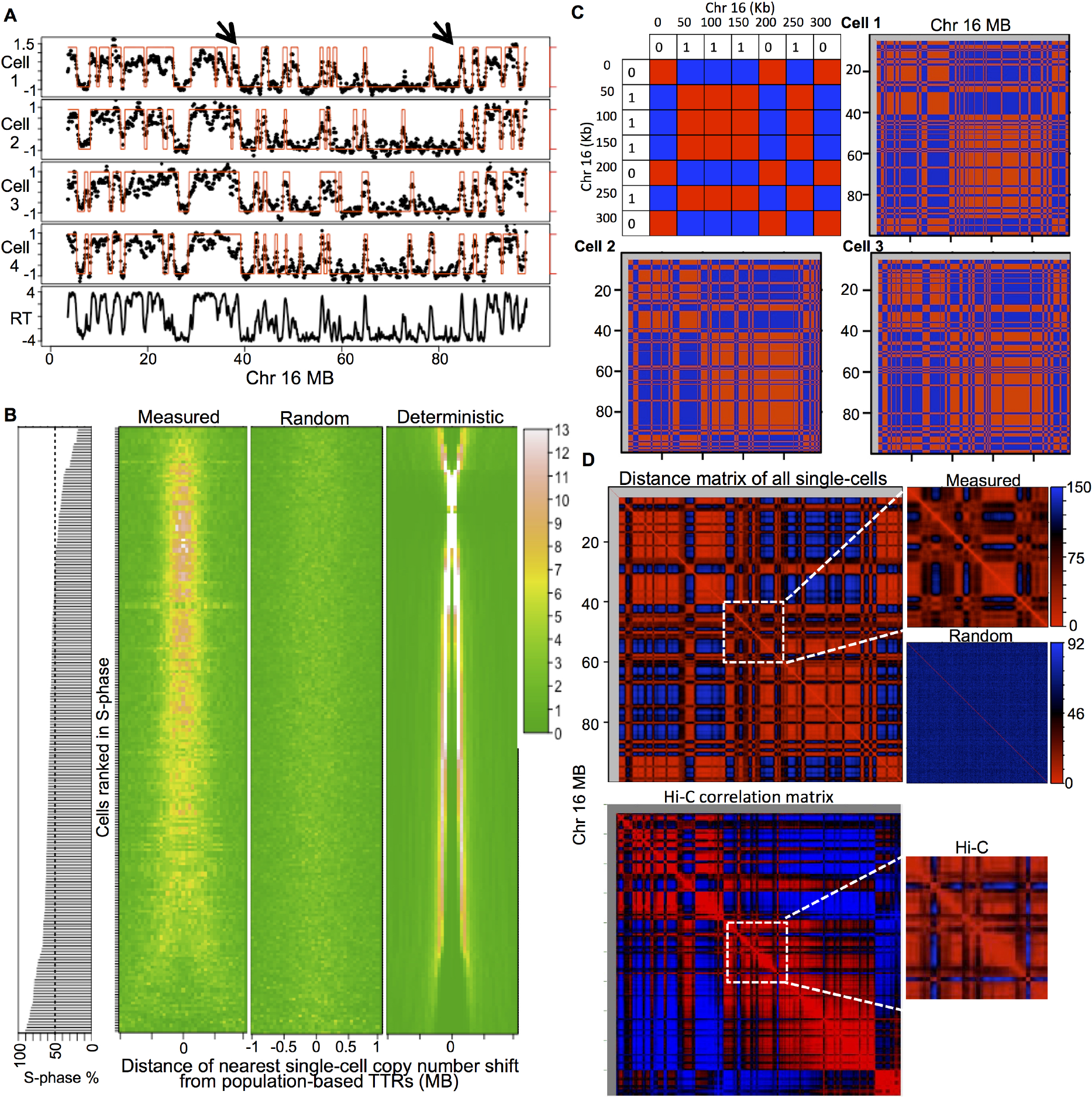
Conservation of TTRs and nuclear compartments in single-cells. a) Binarized single-cell data overlaid on smoothed single-cell CNV profiles. The arrows indicate representative TTRs in population-based data that align with copy number shifts across singlecells. b) Heatmap showing the distance of the closest single-cell copy number shift from the population-based TTR centers. Each row represents one single-cell and the heat map shows percentage of closest single-cell copy number shifts in that cell at different distances from the TTR center. The cells are ranked by position is S-phase. c) Coordinately regulated bins in each single-cells are color coded red and oppositely regulated bins are color coded blue as shown in the top-left schematic. Whole chromosome plot of single-cell from mid-S phase shows coordinated replication across cells and align with Hi-C compartments (shown in panel d). d) Plotting pair-wise absolute distance for all possible pairs of 50 kb bins within each chromosome, using the binary replication status across all cells reveal the functional compartmentalization of replication that show near perfect similarity to Hi-C compartments plotted using Juicebox^28^. The randomized control of single-cell replication lacks compartments highlighted by plotting a 20 Mb segment of Chr 16

In contrast to haploid cells, diploid single-cell replication data can exhibit a third state characterized by asynchrony between homologues where one homologue has replicated and the other remains un-replicated. Since it was not possible to consistently and confidently identify this third state based on copy number alone, we took advantage of hybrid musculus 129 × Castaneus mESCs (subspecies 5mya) that harbor a sufficiently high SNP density to permit parsing of the diploid cell genomes into maternal and paternal haploid genomes. We first segmented the smoothed CNV profiles and binarized them as described for the haploid cell data. Next, we generated homologue specific coverage data by parsing the sequencing reads based on homologue specific SNPs. This allowed us to calculate an *asynchrony score* to measure homologue-to-homologue variability for each segment as the log ratio of 129 to Castaneus read coverage. However, despite the high density of SNPs, homologue parsing nonetheless results in significant loss of coverage, particularly in regions of low SNP density.

Therefore we filtered for those segments with high confidence of replication asynchrony. First we only considered segments with a minimum read number of 40 based on previous empirical determination of read number requirement for efficient copy number identification^18^. Next, the threshold for the *asynchrony score* was set to be outside the distribution of *asynchrony scores* calculated from the control G1 and G2 cells (Supplementary Fig. 4). Homologue specific binarized data was then generated by modifying the original binarized data to reflect segments with asynchrony within each cell (Methods). At the end of allele-specific parsing, 71 diploid S-phase cells had passed all quality control measures and were used for further analysis.

The combined haploid and diploid binarized data ranked by the position of each cell in S-phase (total DNA content) revealed domains with incomplete replication during early S-phase that progressively completed replication in cells ranked later in S-phase (Fig. 2a, Supplementary Fig. 5). These domains correspond to early replicating CTRs identified in population-based replication-timing profiles. Comparison of the binary single-cell replication timing to simulated deterministic (where each cell follows the population-based replication timing accurately) vs. simulated random replication timing indicates a high degree of cell-to-cell replication-timing conservation (Fig. 2a, Methods). Also, the binary replication signal in mid-S cells corresponded to early and late replicating segments in population-based replication timing data (Fig. 2b). Further, since our data reveals that replication timing is well conserved at the single-cell level, we reasoned that in order to detect cell-to-cell variability in replication time for any particular domain, we would need to capture cells at a time during S-phase very close to the average replication time of that domain. In other words, replication-timing variability in early, middle and late S-phase should be maximum for early, mid and late replicating regions, respectively. To test this prediction, we calculated the variability of binarized data for all pair-wise combinations of single-cells that were ranked within one percentile of S-phase progression. Consistent with our prediction, we found a very pronounced increase in replication timing variability at segments whose population-based replication timing corresponded to the position of the single-cell in S-phase (Fig. 2c).

To quantify variability in replication timing during S-phase, we first converted the population-based replication timing in log_2_ enrichment scale to time in hours assuming a 10 hr S-phase^22^ so that each 50 kb bin would have an expected average time of replication in hours. We then converted the percentage progression of each single-cell in S-phase into hours during S-phase. Using these two quantities we generated a cell-specific “time from scheduled replication” profile by subtracting the number of hours the individual cell has progressed in S-phase from the population-based genome-wide replication time in hours for each 50kb bin (Supplementary Fig. 6). Thus for each cell, positive values indicate bins that are scheduled to replicate in the future and negative values indicate bins that should have already replicated.

To estimate the cell-to-cell extrinsic variability in replication timing, we calculated the fraction of cells that replicated each bin position within each possible “time from scheduled replication” in intervals of 0.1 hr. For example, a mid-replicating position can only contribute bins with “time from schedule replication” within +/-5 hr, while a early replicating region that replicates at 2 hours into S-phase, can only contribute bins with a “time from schedule replication” from −8 to +2 hours (total S-phase length is 10 h). Since this calculation was done for all 50 kb bin positions in the genome, we plotted the mean across all bins for each 0.1 h interval of “time from scheduled replication”. Supplementary Figure 7 explains this calculation at an exemplary location in the genome. The kinetics resembled a sigmoid curve that was consistent with previously described theoretical models of stochastic replication-timing regulation based on population-based replication timing data^11,23–25^ (Fig. 2d). To estimate the stochasticity we modeled the kinetics using standard parameters of a sigmoid curve (methods). Then we estimated T_width_, defined as time it takes to progress from 25% to 75% of bins replicated. T_width_ was 2.7 h, which is much lower than the 10 h S-phase, consistent with a stochastic model of DNA replication^11,24^.

To calculate the intrinsic (within cell) replication kinetics, we performed a similar analysis on each haploid cell and homologue-parsed diploid cell independently. We calculated the fraction of bins that were replicated for each possible “time from scheduled replication” in intervals of 0.1 hr across a single genome. The within cell kinetics for all cells was very similar to the cell-to-cell kinetics with an aggregate T_width_ of 2.55 h (Fig. 2e). In contrast, the randomized control had a much larger T_width_ of 15 h and 13.5 h for intrinsic and extrinsic respectively (Fig. 2d, e). Finally, we compared the homologue-to-homologue variation in the diploid cells to “time from scheduled replication”. Homologue-to-homologue variation was measured as percentage of bins that differ between homologues (absolute difference between the binary signal for all homologous pairs of bins). We limited the analysis to 10 cells with the highest read number after homologue specific read parsing. As expected, the maximum variation is at the regions that are actively replicating at the time of harvest (time from scheduled replication = 0 h) (Fig. 2f). The cumulative sum of the variation resembles a sigmoid curve, very similar to the cell-to-cell variation (T_width_=3.1 h). These results demonstrate that extrinsic and intrinsic variation in replication timing are indistinguishable, favoring a model of replication timing regulation where the timing is the outcome of stochastic domain firing and is not affected by the precise environment within a cell. Intriguingly, we could not detect any difference in stochasticity between early and late replicating bins (Fig. 2g), demonstrating that the mechanisms determining the scheduled replication time of any given domain are independent from those determining probability of a domain firing at its scheduled time.

Population-based replication timing data reveals that CTRs are punctuated by segments of DNA that replicate gradually later with time, termed Timing Transition Regions (TTRs)^4^. The binarized single-cell data from mid-S cells reveals conserved locations of copy number shifts that align with TTRs observed in the population-based data (Fig. 3a). The aggregate alignment of TTR centers (mid-transition) to the closest single-cell copy number shifts reveals a strong enrichment of the copy number shifts near the population-based TTR centers (Fig. 3b). In principle, the positions of copy number shifts should show maximum distinction between early and late replicating CTRs in cells that were in mid-S phase. Consistent with this, the alignment of single-cell copy number shifts to TTRs is highest in cells ranked closer to the middle of S-phase (Fig. 3b). The randomized control does not show any enrichment whereas the deterministic simulation shows near perfect enrichment at TTRs in mid-S phase cells. Thus the regions of timing transition from early replication to late replication are conserved between single-cells and population-based data and are also conserved from cell-to-cell.

Chromatin conformation capture studies using Hi-C have revealed the broad segregation of chromatin into two functionally distinct compartments (A and B) that correspond to active and inactive chromatin^26^. This A/B compartmentalization strongly correlates with spatially and temporally distinct early/late replicating CTRs, respectively^5,6^. To measure the prevalence of these compartments in single-cells, we used mid-replicating cells to construct a matrix representing co-regulated bins (pairs of bind that are both replicated or both unreplicated) as red and oppositely regulated (one replicated one not) regions as blue. The co-regulated regions were conserved between cells and recapitulated the nuclear compartmentalization measured by Hi-C with remarkable accuracy (Fig. 3c). Next we calculated a matrix of pair-wise absolute distance for all possible pairs of 50 kb bins within each chromosome, using the binary replication status across all cells. This heat map matrix shows regions of coordinated replication in single-cells and was almost identical to the Hi-C correlation matrix. These results show the conservation of functional (replication timing) nuclear compartments from cell-to-cell that correspond to the structural compartments measured by Hi-C (Fig. 3d).

In summary, we used copy number variation to measure replication timing in single-cells in both haploid and diploid cells. The results support a conserved yet stochastic regulation of replication timing where intrinsic variability and extrinsic cell-to-cell variability are similar, regardless of the timing or chromatin state of each domain. The program is far from random, rather it is highly conserved from cell-to-cell with most individual domains replicating within +/-15% the length of S-phase from their scheduled time. Future studies will be necessary to reveal the molecular events contributing the degree of stochasticity. Here, we have developed methods to measure this degree of stochasticity. We also show that the locations of early to late timing transition observed in population-based replication timing data are conserved in single-cells. It has been previously shown that early replicating and late replicating segments observed in population-based replication timing data align with the Hi-C based A and B sub-nuclear compartments respectively^5,6^. A recent study revealed the conservation of the A and B compartments at the single-cell level^27^. The regions of the genome that are replicated at similar times in single-cells align with these compartments demonstrating their structural and functional conservation. Overall, our results show that that the spatio-temporal DNA replication program is conserved in single-cells, and provides the first direct quantification of single-cell replication timing variability, supporting a stochastic model of replication-timing regulation.

## ACKNOWLEDGMENTS

We thank T. Baslan and J. Hicks for providing indexing barcodes; J. Bechhoefer, B. van Steensel and A. Belmont for their helpful comments. This work was supported by NIH GM083337, GM085354, DK107965 to DMG.

## Author contributions

VD and DMG conceived the study; VD performed the experiments and analysis; VD and DMG wrote the manuscript.

## METHODS

### Cell culture and sorting

Diploid F121-9 hybrid cell line and Haploid H129-2 (ECACC 14040205, a gift from Martin Leeb) mouse ESCs were cultured in feeder free 2i media. To sort single-nuclie, 1 million cells were suspended in 0.5 ml of NST-DAPI buffer. NST buffer was made by mixing the following components in ddH2O for a final volume of 800 ml: 146 nM NaCl, 10 mM Tris base (pH 7.8), 1 mM CaCl2, 21 mM MgCl2, 0.05% (wt/vol) BSA and 0.2% (vol/vol) NP-40. Then 200ml of 106 mM MgCl2 and 10mg of DAPI was added to 800ml of NST buffer to make NST-DAPI buffer. Then we used the FACS AriaII flow cytometer to sort single-nuclie from G1, S or G2 phase into a 96 well plate contain lysis buffer (Sigma SeqPlex, SEQXE-50RXN).

### Population replication-timing profiling

Genome-wide population replication timing was measured using Repli-seq protocol as previously described^21^. Briefly, synchronously cycling cells are pulse labeled with the nucleotide analog 5-bromo-2-deoxyuridine (BrdU) to mark nascent DNA. The cells are sorted into early and late S-phase fractions on the basis of DNA content using flow cytometry. BrdU-labeled DNA from each fraction is immunoprecipitated (BrdU IP), amplified and sequenced. Replication timing is then measured as the log_2_ enrichment of early reads over late reads for each 50 kb bin position across the genome.

### Single-cell sequencing

Single-cell sequencing was performed as described in two previous reports^17,18^. Briefly, the cells were lysed and amplified using the Sigma SeqPlex kit. The WGA products were purified and the WGA universal adaptor sequences were removed as per the kit protocol, leaving NN overhangs. Next unique barcoded illumina adaptors with NN overhangs (gift from Timour Baslan and James Hicks) were ligated to the products from the previous step. Upto 96 uniquely barcoded cells were pooled together and amplified. Samples were then quantified using the Bioanalyzer and qPCR and sequenced on Hiseq 2500 sequencer using 50 bp single-end read format. 11 cells were sequenced without restriction enzyme based WGA universal sequence removal as describe previously^17^. These samples were sequenced at 100bp read length to get sufficient mappable reads after adaptor sequence trimming.

### Read mapping

Reads were demultiplexed based on their unique barcodes. Both cast/129 and H129-2 reads were mapped to mm10 mouse genome assembly using Bowtie 2 with default parameter settings. Reads with MAPQ score of above 10 were retained for further analysis. PCR duplicates were removed using rmdup tool in samtools. Mapped reads were binned into 50 kb windows. Homologue specific read mapping based on SNPs was done using a previously published pipeline^21^.

### Data correction and smoothing

Single-cell sequencing data binned into 50 kb windows were used for the data correction and smoothing. Single-cells with less than 250,000 reads were discarded and reads were then converted to read per million (RPM). Cells with complex karyotype aberrations or complete chromosome loss were discarded while cells with aneuploidy were corrected by normalizing to the mean read density of those chromosomes in the control G1 and G2 cells. Complex karyotype abnormalities which were found to occur in very few cells were identified by plotting the whole genome coverage at 1Mb bins.

G1 and G2 cells showed a uniform coverage profile as expected and thus were used to account for GC bias, whole genome amplification bias and to discard regions with mappability issues. The mean of 5 G1 and cells and 1 G2 cell was calculated for all 50 kb bins. Bins with extreme values (mean RPM>99 percentile and mean RPM< 1 percentile) were identified and masked in all single-cells. To identify repetitive segments of the genome with low mappability, we segmented the mean G1/G2 coverage using Piecewise Constant Fits (PCF) in R using package ‘copynumber’^29^. The parameter used were gamma=3 and kmin=10. Segments with mean RPM lower than 5 percentile were discarded. Finally all single-cells were divided by the mean G1/G2 coverage.

Next each single-cell data was centered and scaled to have an equal inter-quartile range. Extreme values (mean RPM>99 percentile and mean RPM< 1 percentile) in each single data was removed followed by median smoothing with a span of 15 windows.

### Data segmentation and Binarization

The corrected smoothed data was segmented using PCF (R copynumber package) to identify segments with similar copynumber. The parameters used were gamma=3 and kmin=5. Next, the segmented data at 50 kb resolution was used to binarize the domains as replicated or un-replicated. Accurate binarization of the segmented data depends on choosing the correct threshold for each single-cell separately. To this end, we used a brute-force strategy and applied 100 equally spaced thresholds spanning the distribution of the segmented data. At each step the segmented data was binarized and finally the best binary fit was chosen.

The binary fit was calculated using manhattan distance between the binarized data and the un-binarized segmented data. Historically, 1 and 0 are used to denote binary data. However, in order to calculate the similarity based on manhattan distance, the binary values and the segmented data must have similar magnitude. To do this we first used mixture model fitting with 2 components (normalmixEM function in R package mixtools) to identify the model means of replicated and un-replicated fractions in the segmented data. Then the binary values were set as the mean of the components. If the rare occasion when component means were very close (less than 0.7), the binary values were decided based on the skew of the segmented data. This happened predominantly when the cells were in very early or very late S-phase, because the fraction of replicated segments and un-replicated segments respectively were too low for the mixture model to identify as a distinct component. A positive skew (skew>0.2) indicated cells that are in early S-phase and the binary values were set to 50 percentile and 95 percentile of the segmented data. A negative skew (skew < −0.2) indicated cells that are in late S-phase and the binary values were set to 5 percentile and 50 percentile of the segmented data. Otherwise, the binary values were set to 25 percentile and 75 percentile of the segmented data. The binary signal with the minimum manhattan distance (highest similarity) from the segmented data was chosen as the best fit (**Supplementary movie 1**). Outlier bins with a segmented value outside of +/-2 were not used for the threshold calculation.

### Homologue specific binary signal

Homologue specific binary signal for diploid hybrid cells were generated based on the homologue parsed sequencing data. First, we used un-parsed data to first segment and then binarize as described above. Two, identical copies of the binary signal were generated and assigned one to 129 allele and the other to the Castaneus allele. The binary signals from each allele can then be modified for segments that are identified to be asynchronous. To identify asynchronous segments, we calculated an *asynchrony score* (R) for each segment as:

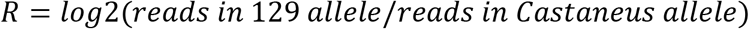

Theoretically, a domain that is completely replicated in one homologue and completely un-replicated in the other homologue will yield an R score of 1 or −1. However, this ideal value cannot be expected due to two reasons. 1) Due to the sparse nature of single-cell data, the large domains identified as replicated or un-replicated by the segmentation and binarization may still contain small segments that differ in their replication status. 2) The distribution of SNPs is non-homogenous resulting in loss of coverage after parsing.

Therefore we discarded segments with a total read number is less than 40. This number was determined based on a previous study that empirically determined total number of reads required for efficient copy number analysis^18^. Next, the threshold for the *asynchrony score* was set to be outside the distribution of *asynchrony scores* calculated from the control G1 and G2 cells. The thresholds used were R> 1.11 for 129 allele to be classified as replicated and R< −1.18 for the Castaneus allele to be classified as replicated.

### Removing outlier cells

Outlier cells were defined as cells that don’t correlate with any other cells after binarization. A heat map of genome-wide manhattan distance using binarized data for every pair-wise combination of haploid cell ordered by their rank in S-phase reveals cells that are ranked close together have the least distance and cells ranked far apart in S-phase have maximum distance. But some cells have low similarity to all the other cells and appear as streaks in the heat map (Supplementary Fig. 3). These cells were removed from further analysis.

### Calculation of Deterministic and Randomized model

The deterministic model assumes that each cell follows the population-based replication-timing profile precisely. For a given single-cell with ‘x’ percentage of the genome replicated, the deterministic profile is given my assigning all 50 kb bins with average population-based timing greater ‘x’ percentile as replicated. To generate the random profile, ‘x’ percentage of 50 kb bins selected randomly are assigned as replicated.

### Calculating sigmoid fit

The sigmoid fit was modeled using a non-linear least squares approach (function *nls* in R) using the formula:

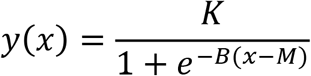

where, K is the maximum value of the sigmoid, B is the growth rate and M is the x intercept at the sigmoid’s mid-point. The initial values of K, B and M were set to 100, 0.7 and 0.

### SUPPLEMENTARY FIGURES

**Supplementary Figure 1:**
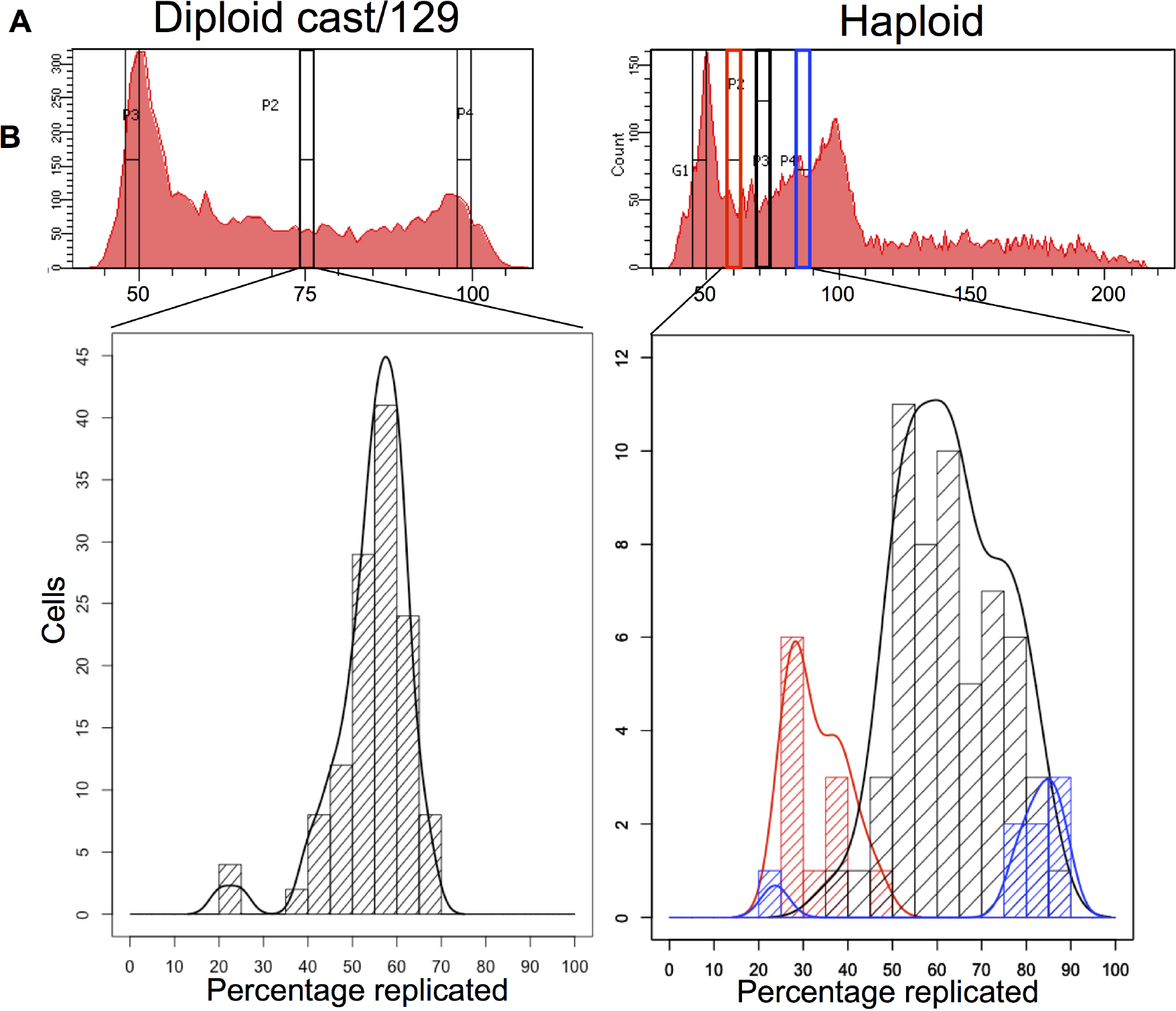
Single-cell sequencing ranks cells accurately in S-phase. a) FACS (Fluorescence Activated Cell Sorting) profile based on DNA content and sorting gates used for diploid and haploid cells. Mid-S-phase cells were sorted for diploid cells. For haploid cells, 12 early S-phase cells (red gate), 12 late S-phase cells (blue gate) and 62 mid S-phase cells (black gate) were sorted. b) Ranking the cells in S-phase after binarization accurately reflects the position of cells in S-phase in both haploid and diploid cells. For reasons we don’t understand, the FACS profile of haploid cells show a high proportion of late S-phase cells. Thus, while majority of the cells sorted using the mid-S-phase FACS gate (black gate) were middle replicating cells, there were also cells from later stages in S-phase. This is accurately reflected by the haploid S-phase rank histogram which shows a skew towards late S-phase. The skew was inconsequential and in fact provided a wider distribution of S-phase cells for subsequent analyses.

**Supplementary Figure 2:**
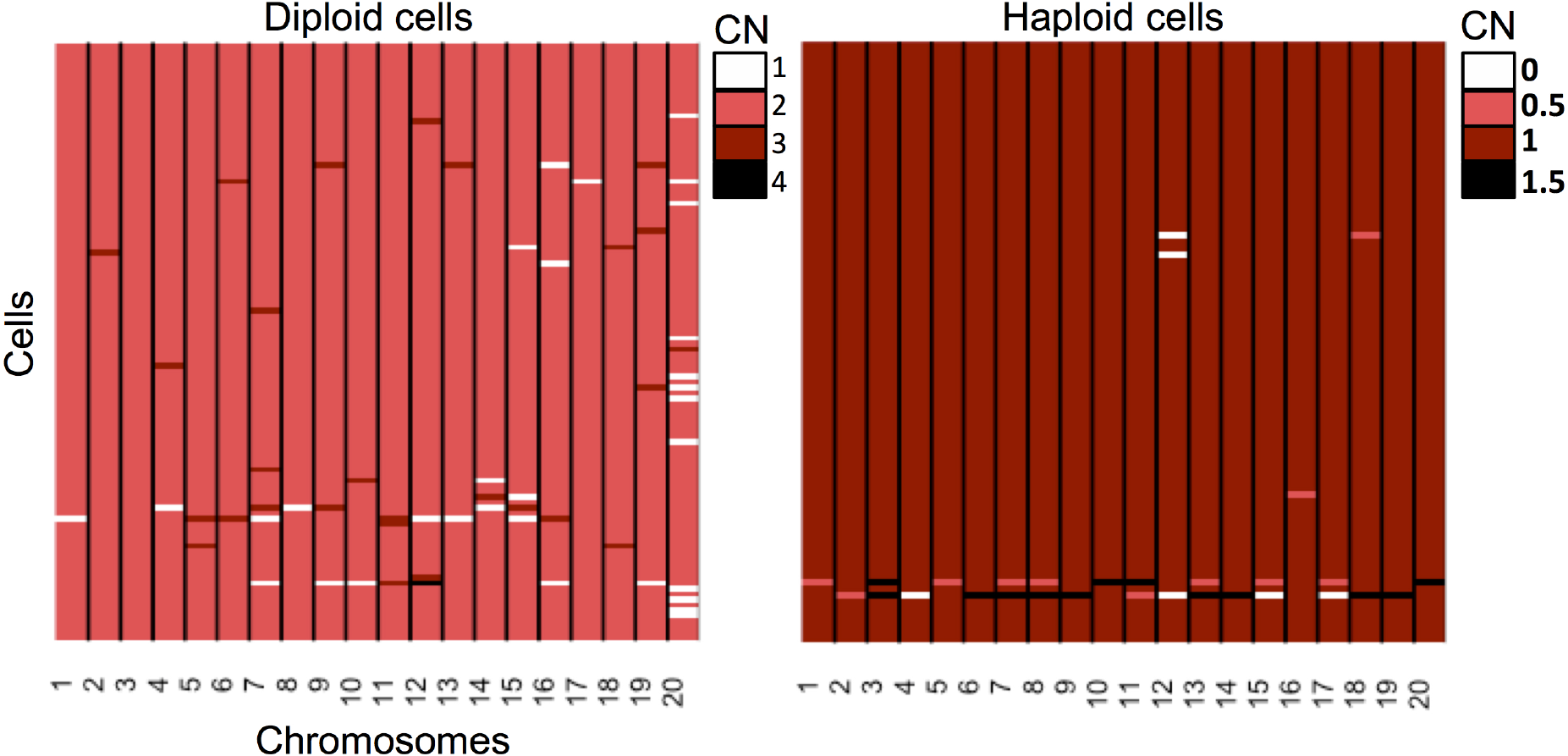
Karyotype aberrations in diploid and haploid cells. Heat map of whole chromosome karyotype aberrations. Approximately 27 percentage of diploid cells show karyotype aberrations. The intermediate copy number observed in 4 haploid cells may be due to random diploidization.

**Supplementary Figure 3:**
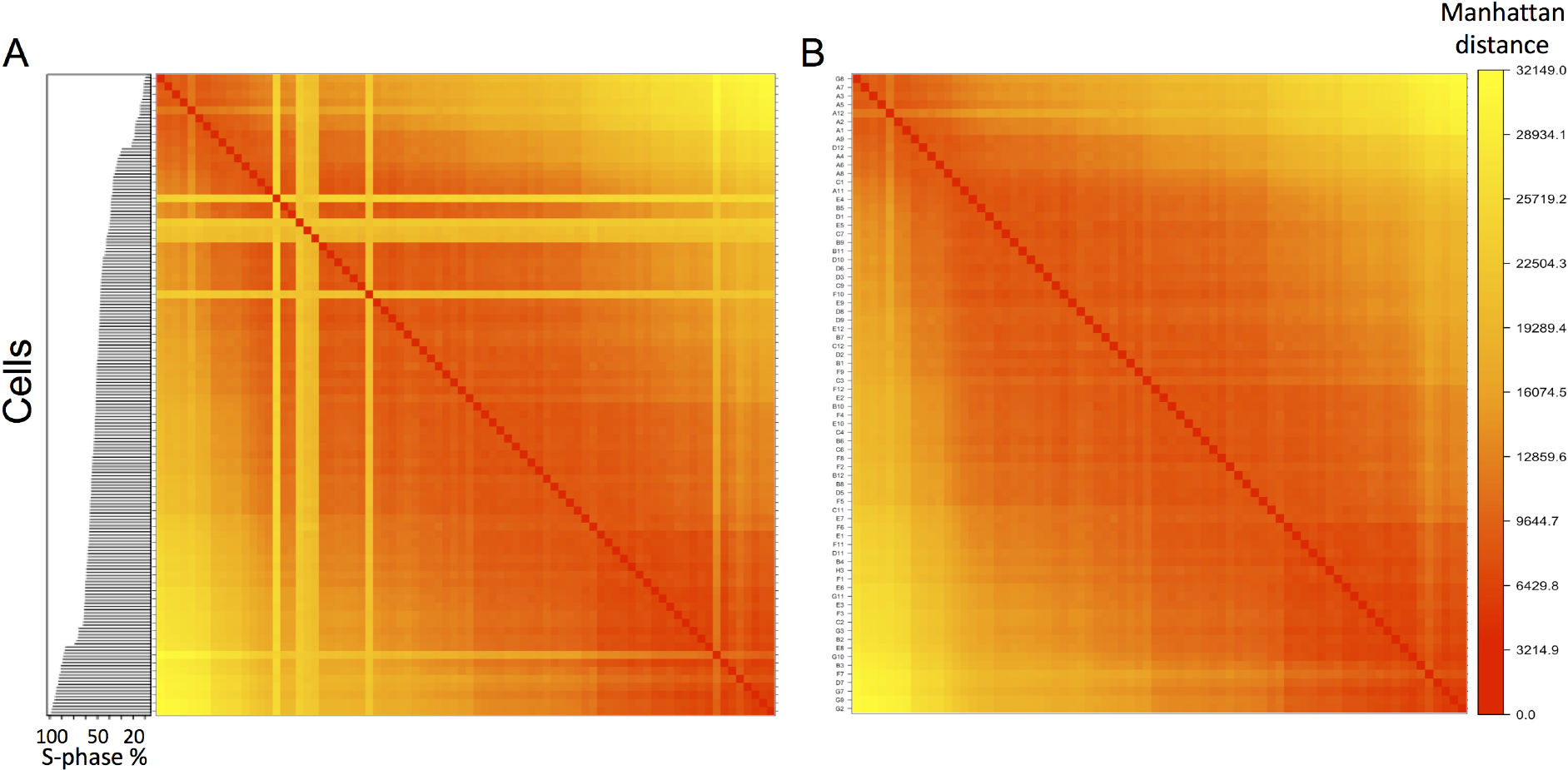
Removing outlier cells. a) Heat map of genome-wide manhattan distance using binarized data for every pair-wise combination of haploid cell ordered by their rank in S-phase. The ranking is shown as a barplot on the left. Outlier cells that do not correlate with any other cells appear as streaks on the heat map. b) Heat map after removing the outlier cells.

**Supplementary Figure 4:**
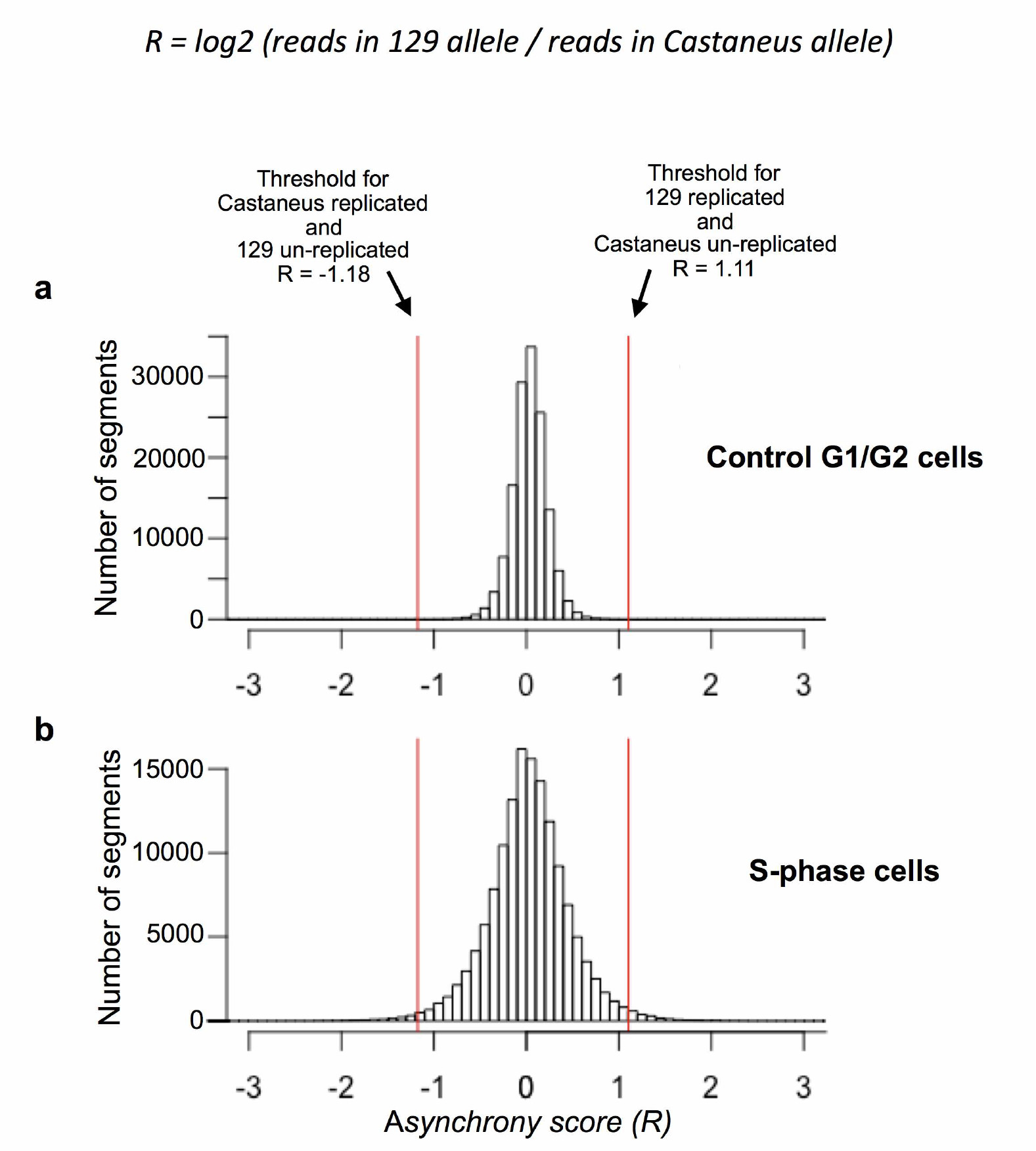
Calculating the threshold asynchrony score to define segments in diploid cells that show homologue asynchrony. a) Histogram of asynchrony score (R) calculated using the control G1/G2 cells. b) Histogram of asynchrony score in S-phase cells. The thresholds (red lines) were set to be the maximum and minimum asynchrony score in the control G1/G2 cells.

**Supplementary Figure 5:**
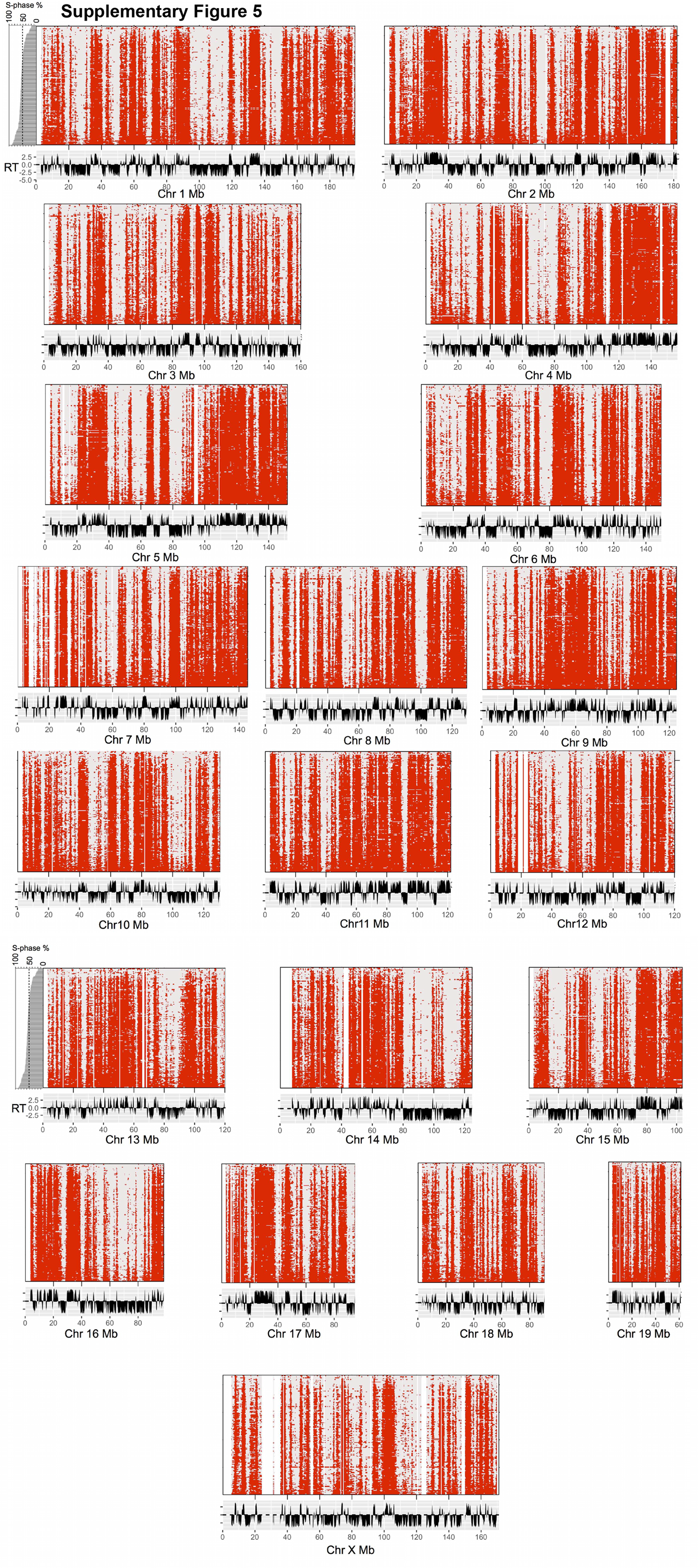
Binararized data ranked by progress in S-phase for 75 haploid cells and 71 diploid diploid parsed into paternal and maternal genomes for all chromosomes.

**Supplementary Figure 6:**
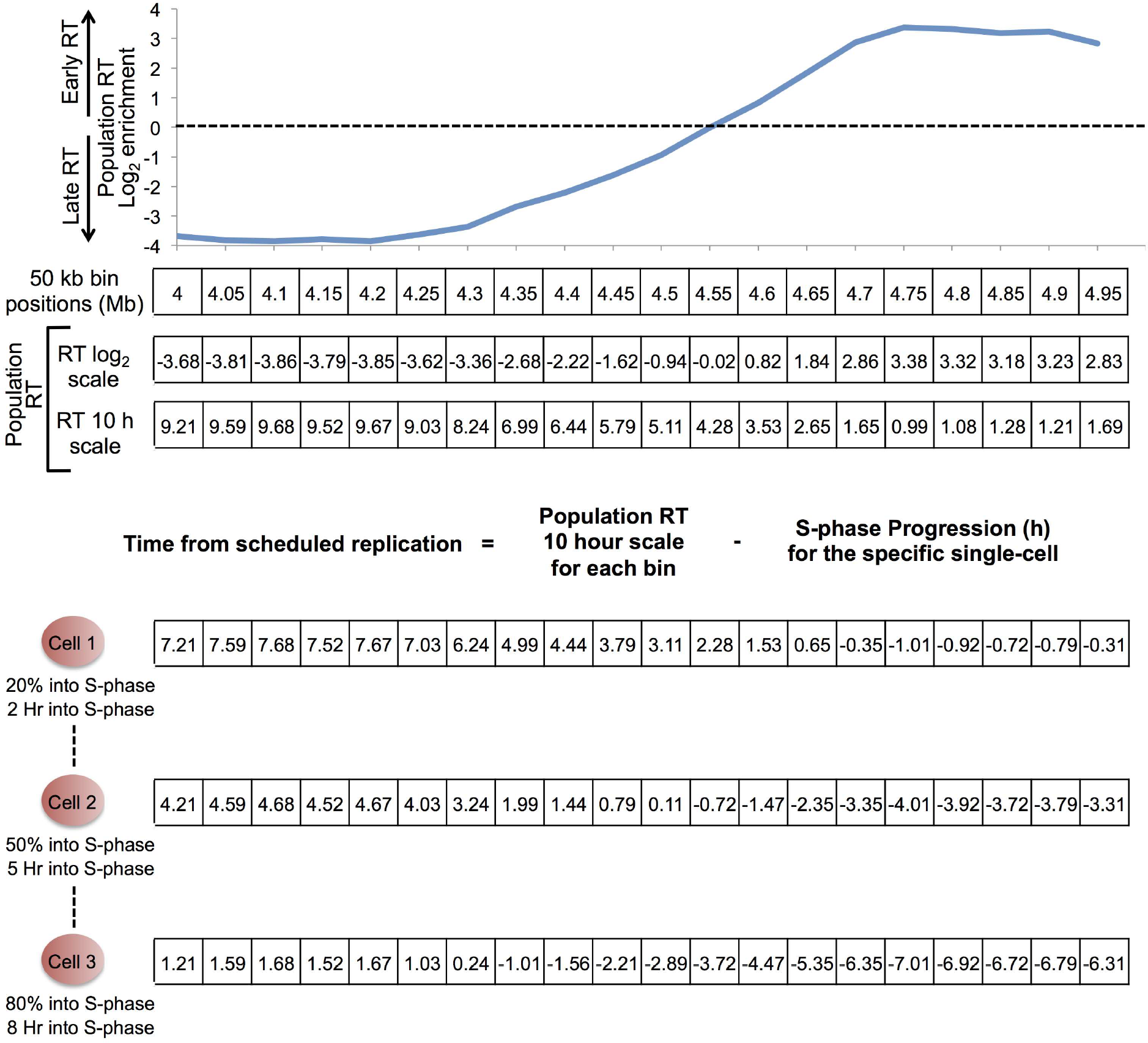
Generation cell-specific “Time from scheduled replication”. The steps involved in the generation of cell-specific “Time from scheduled replication” is shown for a 1 Mb example region. The population replication-timing values in log2 enrichment scale was converted to a 0-10 hour scale such that each 50 kb bin position has a population-based replication-timing value in hours. In this scale, the late replicating bin positions will have values close to 10 h and early replicating bin positions will have values closer to 0 h. Next for each cell, it’s progression in S-phase was converted to hours. The cell-specific “time from scheduled replication” profile was generated by subtracting the number of hours the individual cell has progressed in S-phase from the population-based genome-wide replication time in hours for each 50 kb bin. Thus for each cell, positive values indicate bins that are scheduled to replicate in the future and negative values indicate bins that should have already replicated.

**Supplementary Figure 7:**
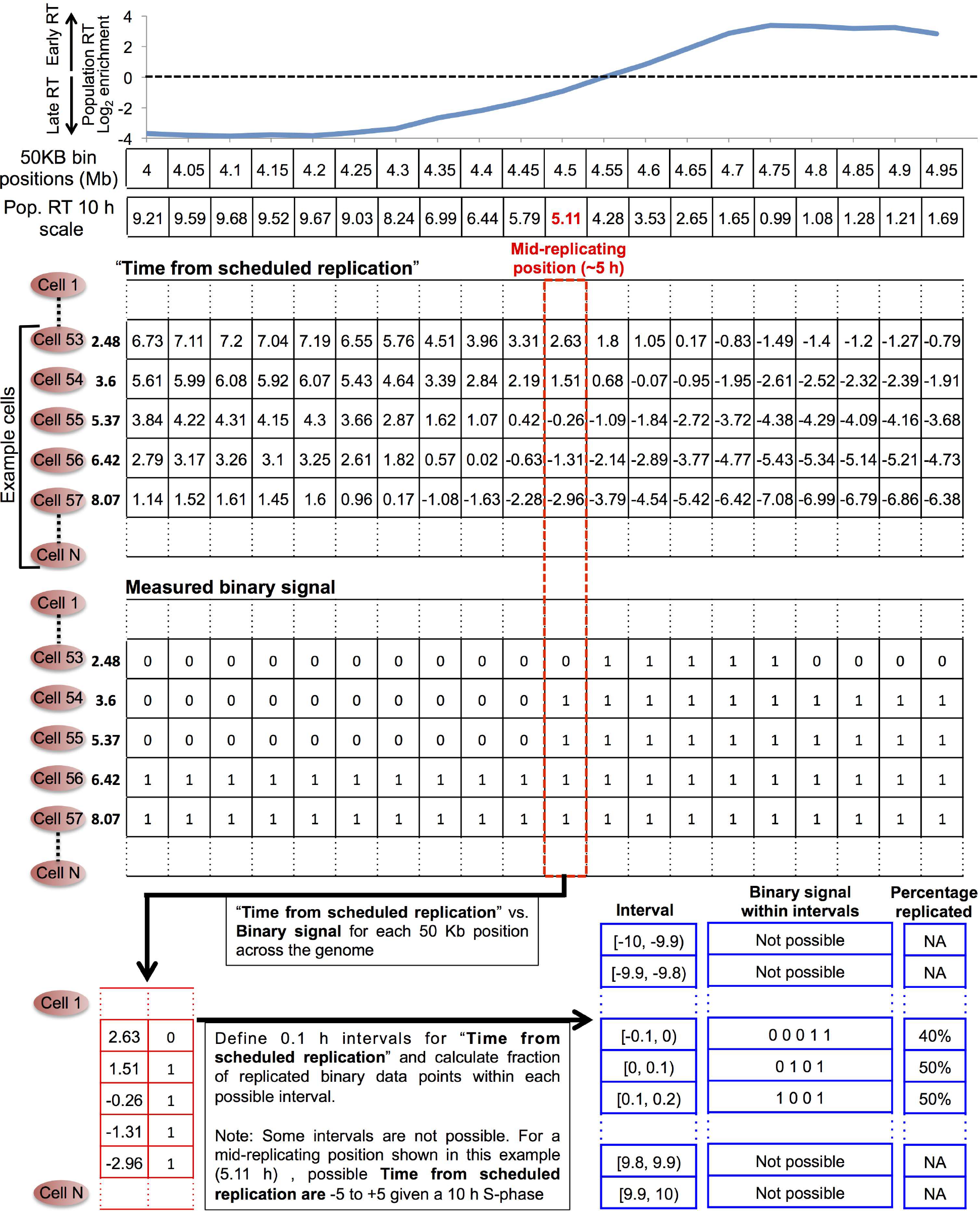
Cell-to-cell extrinsic variability calculation. The goal of this analysis is to compare the cell-to-cell variability as a function of the “Time from scheduled replication”. The example here shows this calculation for a mid-replicating 50 kb bin position. The “Time from scheduled replication” and the measured binary signals are collected for this bin position across all cells and the data is split into 0.1 h intervals of “Time from scheduled replication”. The fraction of replicated data points is calculated for each interval. This analysis was repeated for all bin positions across the genome. Then the mean across all bin positions were plotted for each 0.1 h interval of the “Time from scheduled replication” (Fig. 2d).

